# Genetic characterization of *Streptococcus equi* subspecies *zooepidemicus* associated with high swine mortality in United States

**DOI:** 10.1101/2019.12.12.874644

**Authors:** Xuhua Chen, Nubia Resende-De-Macedo, Panchan Sitthicharoenchai, Orhan Sahin, Eric Burrough, Maria Clavijo, Rachel Derscheid, Kent Schwartz, Kristina Lantz, Suelee Robbe-Austerman, Rodger Main, Ganwu Li

**Affiliations:** Department of Veterinary Diagnostic and Production Animal Medicine, College of Veterinary Medicine, Iowa State University, 1800 Christensen Drive, Ames, IA 50011, USA; National Veterinary Services Laboratories, 1920 Dayton Ave, Ames, IA 50010

## Abstract

High mortality events due to *Streptococcus equi* subspecies *zooepidemicus* (*S. zooepidemicus*) in swine have not previously been reported in the United States. In September and October 2019, outbreaks with swine mortality up to 50% due to *S. zooepidemicus* septicemia were reported in Ohio and Tennessee. Genomic epidemiological analysis revealed that the eight outbreak isolates were clustered together with ATCC 36246, a Chinese strain caused outbreaks with high mortality, also closely related to three isolates from human cases from Virginia, but significantly different from an outbreak-unrelated swine isolate from Arizona and most isolates from other animal species. Comparative genomic analysis on two outbreak isolates and another outbreak-unrelated isolate identified several genomic islands and virulence genes specifically in the outbreak isolates only, which are likely associated with the high mortality observed in the swine population. These findings have implications for understanding, tracking, and possibly preventing diseases caused by *S. zooepidemicus* in swine.

## Introduction

*Streptococcus equi* subspecies *zooepidemicus* (*S. zooepidemicus*), a beta-hemolytic and Gram-positive bacterium, is most frequently isolated as an opportunistic pathogen of horses in the upper respiratory and lower genital tracts. It can also cause infections in a wide range of other animal species, including cats, ruminants, pigs, monkeys, dogs, and guinea pigs (*1–6*). *S. zooepidemicus* is zoonotic, with reported transmission from horses, dogs, and guinea pigs to humans (*7*) leading to either severe invasive disease (bacteremia, septic arthritis, pneumonia, and meningitis) or benign disease like pharyngitis, with potential to trigger acute post-streptococcal glomerulonephritis (APSGN) (*8*). The human patients usually acquire the bacteria through direct contact with infected animals or consumption of contaminated animal products such as milk or cheese.

The pathogenesis of *S. zooepidemicus* is not fully understood; however, virulence factors and the host immune status appear to have roles in the development of disease. The M protein, a member of M/M-like protein family, is a surface-associated protein and a classical virulence factor in group A *Streptococcus* (GAS; e.g., *S. pyogenes*) (*9*). The M-like protein SzP of *S. zooepidemicus* was reported to contribute to the virulence in animal model studies (*10,11*) and a second M-like protein SzM has been shown to bind fibrinogen, activate plasminogen, inhibit phagocytosis, and serve as a protective antigen for vaccination (*12*). Very recently, BifA, a Fic domain-containing protein was shown to disrupt the blood-brain barrier integrity by activating moesin in endothelial cells (*13*). In addition, capsular and other surface polysaccharides are important streptococcal virulence factors, several superantigen genes (*seeM, szeL,* and *szeM*) and several genes associated with anti-phagocytic capability in *S. zooepidemicus* may be involved in the pathogenicity (*14*).

Although *S. zooepidemicus* was reported to cause epizootic outbreaks in swine resulting in significant economic losses in China and Indonesia (*4, 15*), previous isolation of *S. zooepidemicus* from clinically ill pigs have been rather limited in the US in past decades. The first high mortality event from *S. zooepidemicus* in North America was reported very recently in Canada in March 2019 (https://www.biorxiv.org/content/10.1101/812636v2). From late September to early October of 2019, three cases of cull sows and feeder pigs from Ohio and Tennessee were submitted to the Veterinary Diagnostic Laboratory at Iowa State University (ISU-VDL). High mortality ranging from 10-50% in groups of pigs was reported over the period of 8-10 days at the buying station in Ohio and similar high mortality (922 out of 2,222 sows in lairage) from an abattoir in Tennessee. The clinical signs included sudden death, weakness, lethargy, and high fever. Splenomegaly and hemorrhagic lymph nodes were the most consistent macroscopic findings. Microscopic lesions were consistent with acute bacterial septicemia. A laboratory diagnosis of *S. zooepidemicus* septicemia was given, which was corroborated by histopathology, PCR, and bacterial culture. To genetically characterize *S. zooepidemicus* strains associated with high mortality and gain insights into the epidemiology of these highly unusual and unexpected outbreaks, we performed whole genome sequencing on eight isolates from the Ohio and Tennessee outbreaks, another outbreak-unrelated swine isolate from Arizona, and 15 *S. zooepidemicus* isolates from other animal species. Three full-length complete genome sequences were further assembled and genomic epidemiological and comparative genomic analyses were conducted.

## Materials and Methods

### *S. zooepidemicus* isolates

In total, twenty-four *S. zooepidemicus* isolates were whole genome sequenced and included in this bacterial genomic epidemiological study. Among them, eight isolates (OH-71905, TN-74097, NVSLTN-LUNG1, NVSLTN-LUNG2, NVSLTN-LUNG3, NVSLTN-LIVER4, NVSLTN-TB1 and NVSLTN-TC1) were from Ohio and Tennessee outbreaks, another swine isolate (AZ-45470) from a case not related to the outbreaks, and 15 isolates from different animal species (6 from equine, 3 from feline, 3 from guinea pig, 1 from canine, 1 from caprine, and 1 from chinchilla). Another 24 strains from different countries and years with complete or draft genomes that were publicly available from GenBank (https://www.ncbi.nlm.nih.gov/genbank/) were included in study. The detailed information of all 48 *S. zooepidemicus* strains are summarized in Appendix Table 1-2.

### DNA extraction and library preparation

A single colony of each *S. zooepidemicus* strain was inoculated into Tryptic Soy Broth (TSB) with 10% bovine serum and incubated at 37°C with overnight shaking. ChargeSwitch gDNA mini bacteria kit (Life Technologies, Carlsbad, CA) was used to extract genomic DNA from *S. zooepidemicus* cells following the manufacturer’s guidelines. DNA quality was determined by NanoDrop (Thermo Fisher Scientific) and accurate concentration was measured by Qubit fluorometer double-stranded DNA high sensitivity (dsDNA HS) kit (Life Technologies). At the National Veterinary Service Laboratories (NVSL), a loop of *S. zooepidemicus* from an overnight culture grown on blood agar was inoculated into 400 microliters of TE buffer containing 2.5 mg of lysozyme and incubated for 2 hours at 37°C. The entire volume was extracted using a Promega Maxwell RSC Whole Blood DNA Kit on the Promega Maxwell RSC 48 (Promega, Madison, WI). Accurate DNA concentration was measured by Qubit fluorometer double-stranded DNA high sensitivity (dsDNA HS) kit (Life Technologies). Indexed genomic libraries were prepared at both laboratories by using Nextera XT DNA library prep kit (Life Technologies) for subsequent sequencing.

### Genome sequencing and assembly

Bacterial genomes were sequenced using the Illumina MiSeq platform (Illumina) with 250×2 read length in the NGS Unit of the ISU-VDL or at the NVSL. Low-quality raw reads and adapters were filtered and trimmed by Seqtk and Trimmomatic-0.36 (*16*). The filtered reads were tested for quality by FastQC and were assembled utilizing SPAdes 3.13.1-Darwin (*17*). Assembly quality was evaluated using Quast (*18*) to determine N50, longest contigs, total length of contigs, GC content, and other parameters as appropriate. Contigs with low average depth (≤ 2) or small coverage (≤ 500) were removed from further genomic analysis. The genome sequencing and assembly data were deposited at NCBI under BioProject accessions PRJNA588803 (VDL) and PRJNA591128 (NVSL).

### Phylogenetic analysis

Single nucleotide polymorphisms (SNPs) of 48 *S. zooepidemicus* isolates were identified by running kSNP3 with standard mode. The optimal k-mers size was calculated by Kchooser program and the whole-genome phylogeny was analyzed based on identified core genome SNPs (*19*). Sequence type (ST) based on 7 highly conserved housekeeping genes (*arc, nrdE, proS, spi, tdk, tpi* and *yqiL*) was assigned for each *S. zooepidemicus* genome according to the PubMLST *S. zooepidemicus* database (http://pubmlst.org/szooepidemicus)(*20*). Tree Of Life (iTOL, https://itol.embl.de/) (*21*) was used for display, manipulation and annotation on the base of core SNPs tree.

### Genome gap closure and comparative genomic analysis

Three genomes of *S. zooepidemicus* including two outbreak strains of OH-71905 from Ohio and TN-74097 from Tennessee, and another outbreak-unrelated swine isolate AZ-45470 from Arizona were further sequenced using Oxford Nanopore Technologies (Oxford, United Kingdom) GridIONx5 in the DNA facility at ISU to generate longer reads for genome gap closure. The full-length circular genome sequences were obtained by hybrid assembly combining both Illumina short reads and Nanopore long reads using Unicycler v0.4.8 (*22*). Three complete genome sequences are available at NCBI under BioProject accession PRJNA588803. Comparative genomic studies were performed with BLAST Ring Image Generator (BRIG)(*23*) to generate a circular genomic map using three closed genome sequences in addition to two complete sequences of control strains ATCC 35246 and CY (Nanjing) strains (GenBank accession number CP002904.1 and CP006770.1). The circular graphical map was plotted to feature GC skew, GC content, and predicted genomic islands (GIs) along with genome comparisons.

### Virulence gene and genomic islands identification

Putative virulence genes were retrieved from genome sequences according to previous publications (*13, 24–27*). The prediction of genomic island (GI) was based on IslandPath-DIMOB, SIGI-HMM, IslandPick and Islander using IslandViewer 4 (*28*). Coding sequences (CDS) in every GI were annotated by NCBI Prokaryotic Genome Annotation Pipeline (PGAP) (*29*). The distribution of putative virulence genes and proportion of CDS of GIs in all of the *S. zooepidemicus* isolates were determined in R using pheatmap package. All gene sequences including putative virulence genes and CDSs in GIs were identified by local Blast+ (2.9.0 version) choosing BLASTn option.

## Results

### Phylogenetic characterization of *S. zooepidemicus*

Whole genome sequencing was performed on 24 *S. zooepidemicus* isolates. More than 90% of the paired-end reads processed had a Phred score of 35, indicating the high quality of sequencing data. The *de novo* assembly of raw reads generated contigs with total size ranging from 2.0 Mbp to 2.2 Mbp, and an overall GC content between 40% and 43%, which were similar to those of reference genome ATCC 35246 (2,167,164bp, 41.65% GC). Thus, the quality of all assembled contigs of 24 isolates were considered sufficient for whole genome analysis.

Whole genome phylogenetic analysis based on core genome SNPs was conducted with 48 *S. zooepidemicus* isolates including the 24 isolates sequenced in this study and 24 strains from different countries with publically available genome sequences (Figure 1). A total of 23,659 core genome SNPs were identified from all strains by kSNP3. The phylogenetic tree based on the core genome SNPs indicated a large genetic diversity and the 48 isolates could be clustered into 33 phylogenetic lineages (threshold = 0.01) with only 5 clusters consisting of more than one strain (isolate), while the other 28 clusters contained only one strain (isolate). No obvious geographic or historic distribution differences could be found in our phylogenetic analysis. The eight *S. zooepidemicus* isolates from Ohio and Tennessee outbreaks were indistinguishable based on whole genome phylogeny, suggesting that they shared a common source. Surprisingly, the eight outbreak isolates were clustered together with the Chinese strain (ATCC 36246), which caused outbreaks of high mortality in 1975 (*15*). In addition, three isolates from human cases with guinea pig exposure (NVSLVA-S19, NVSLVA-S2, and NVSLVA-S22) (*6*) were also closely related to the outbreak isolates. In contrast, another swine isolate (AZ-45470), which did not cause a high mortality event, was distant to the outbreak isolates.

**Figure 1.**
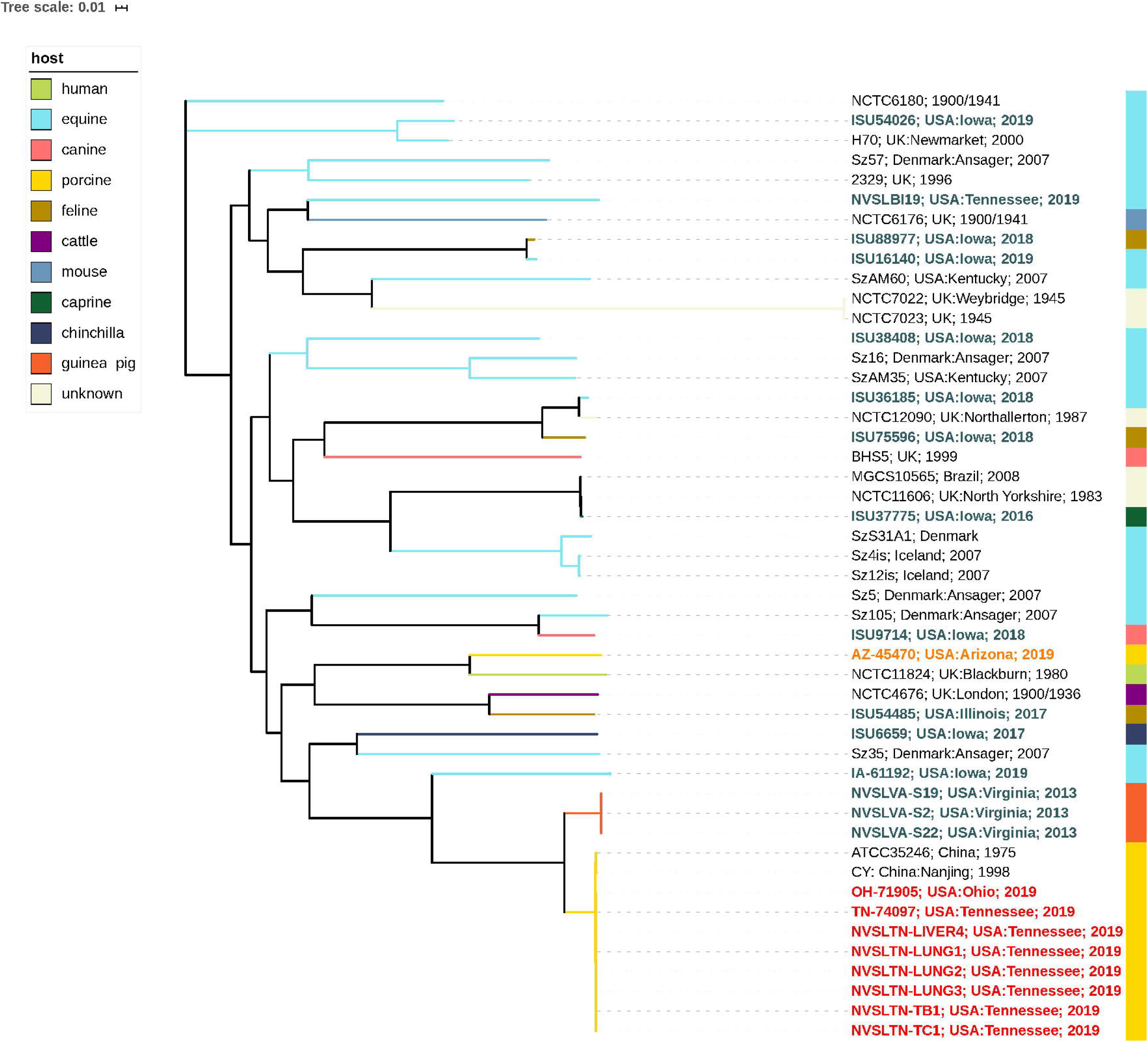
Whole-genome sequence-based phylogenetic analysis was conducted using SNPs located in all tested *S. zooepidemicus* genome to generate a core SNP parsimony tree. The branches of the tree are proportional to the distance between the isolates. 11 color strips demonstrate different hosts of total 48 isolates.

The MLST analysis on all 48 isolates revealed 24 previously described ST types, while 11 strains (22.91%) represented by 9 novel allelic profiles did not match to any STs available in the current PubMLST database of *S. zooepidemicus* as of December 2019 (Figure 2 and Appendix Table 3). Even though clusterings based on whole genome core SNPs and MLST allelic profiles were overall similar, strains with the same MLST types were distributed both within the same and adjacent whole genome core SNP clusters in the phylogenetic tree, indicating that the SNP-based genotyping was more discriminatory, as expected. The eight strains from Ohio and Tennessee outbreaks and two strains (CY and ATCC 36246) from China were grouped together in ST 194, while the outbreak-unrelated Arizona swine isolate (AZ-45470) was clustered distantly in ST340. Three guinea pig isolates from human cases (NVSLVA-S19, NVSLVA-S2, and NVSLVA-S22) with the same allelic profile and an unassigned ST type, differing from ST 194 at two allele sequences, were closely clustered with the eight outbreak strains from Ohio and Tennessee as well as the Chinese outbreak strain ATCC 36246 within the ST194 lineage.

**Figure 2.**
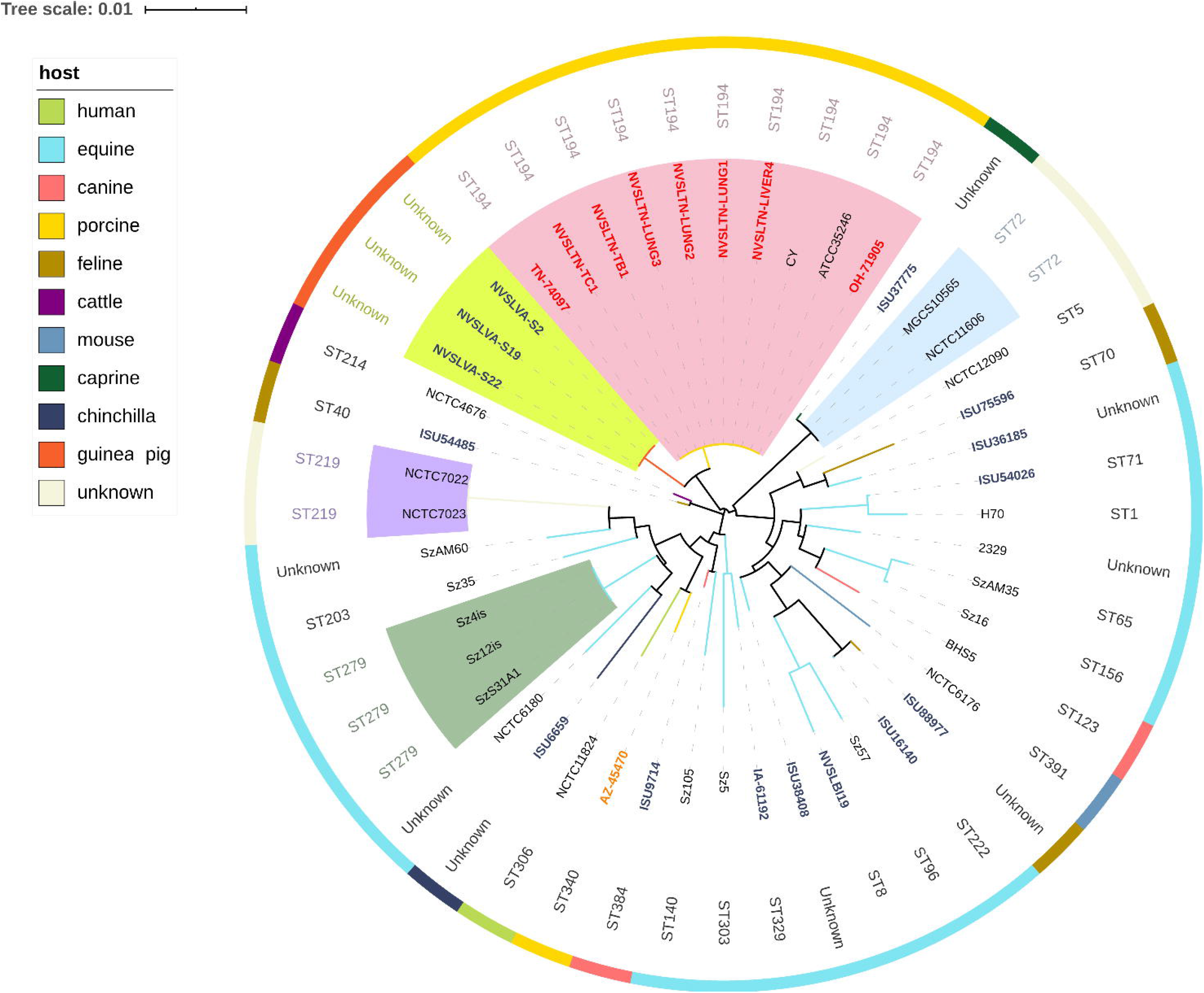
MLST analysis of 48 *S. zooepidemicus* isolates. The outer color ring represents different hosts. ST type of every isolate is listed at the inner circle. The neighbor-joining tree was based on a concatenated alignment of 7 housekeeping genes. Color shades indicate different isolates with identical ST type.

### Comparative genomic analysis of *S. zooepidemicus*

To further characterize the *S. zooepidemicus* isolates associated with the high mortality outbreaks in Ohio and Tennessee (suggesting a hypervirulence trait), the genome gaps of three isolates were closed to obtain full-length genome sequences and perform comparative genomic analysis. Considering the indistinguishability of seven porcine isolates from Tennessee, one representative isolate from Tennessee (TN-74097), the single swine outbreak isolate from Ohio (OH-71905), and the outbreak-unrelated swine isolate from Arizona (AZ-45470) were subjected to Oxford Nanopore Sequencing to generate long reads. Three full-length complete genome sequences (Accession numbers: CP046040, CP046042, and CP046041) were obtained using hybrid assembly of both Illumina short reads and Nanopore long reads. The general genomic features of all three genomes are summarized in Table 1. Both *S. zooepidemicus* strains OH-71905 and TN-74097 contained a circular chromosome of 2,189,155 bp with an average GC content of 41.65%. There was only one base pair nucleotide difference detected in these two outbreak isolates. Genome Annotation Pipeline (PGAP) detected 1,994 CDSs, 67 tRNA genes, 18 rRNA genes, and 4 ncRNA genes in both isolates OH-71905 and TN-74097. Compared to two outbreak isolates from Ohio and Tennessee, the genome size of *S. zooepidemicus* strain AZ-45470 from an outbreak-unrelated case was noticeably smaller with 2,074,453 bp of length and an average GC content of 41.54%. Accordingly, 1,869 CDSs were identified in the genome of AZ-45470, although it had the same numbers of tRNA, rRNA, and ncRNA genes as with OH-71905 and TN-74097 genomes.

**Table 1.** General genomic features of three swine *S. zooepidemicus* isolates OH-71905, TN-714097 and AZ-45470.

| Feature | OH-71905 | TN-714097 | AZ-45470 |
| --- | --- | --- | --- |
| Length, bp | 2,189,155 | 2,189,155 | 2,074,453 |
| GC Content | 41.65% | 41.65% | 41.54% |
| Genes (total) | 2,083 | 2,083 | 1,958 |
| CDSs (total) | 1,994 | 1,994 | 1,869 |
| rRNAs | 6, 6, 6 (5S, 16S, 23S) | 6, 6, 6 (5S, 16S, 23S) | 6, 6, 6 (5S, 16S, 23S) |
| tRNAs | 67 | 67 | 67 |
| ncRNAs | 4 | 4 | 4 |
| Pseudo Genes (total) | 75 | 75 | 47 |

\*CDS, coding sequence; tRNA, transfer RNA; rRNA, ribosomal RNA; ncRNA, non-coding RNA.

Comparative genomic analysis was performed with OH-71905, TN-74097, AZ-45470 genomes as well as the genome of ATCC 35246 and CY strains from China (Figure 3). Genome differences were visualized using BLAST Ring Image Generator (BRIG). As previously mentioned, the genome sequences of OH-71905 and TN-74097 were nearly identical except one bp nucleotide (a single SNP) difference. Overall, OH-71905 and TN-74097 showed an average nucleotide identity of 99.96% with ATCC 35246 within the 99.36% aligned sequence length, indicating that they are highly similar. Compared with the ATCC 35246 genome, 18 insertions and 6 deletions were found in the genomes of OH-71905 and TN-74097, and the largest insertion was 12,494 bp of length from the position 786,295 bp to position 798,785 bp. In contrast, the genome sequence of AZ-45470 displayed an average nucleotide identity of 97.20% with those of OH-71905 and TN-74097 and covering 91.12% of aligned sequence length. Additionally, 189 deletions and 186 insertions were identified in the genome sequence of AZ-4570 compared with those of OH-71905 and TN-74097. The top six largest deletions ranged from 5,000 bp to 50,000 bp in length.

**Figure 3.**
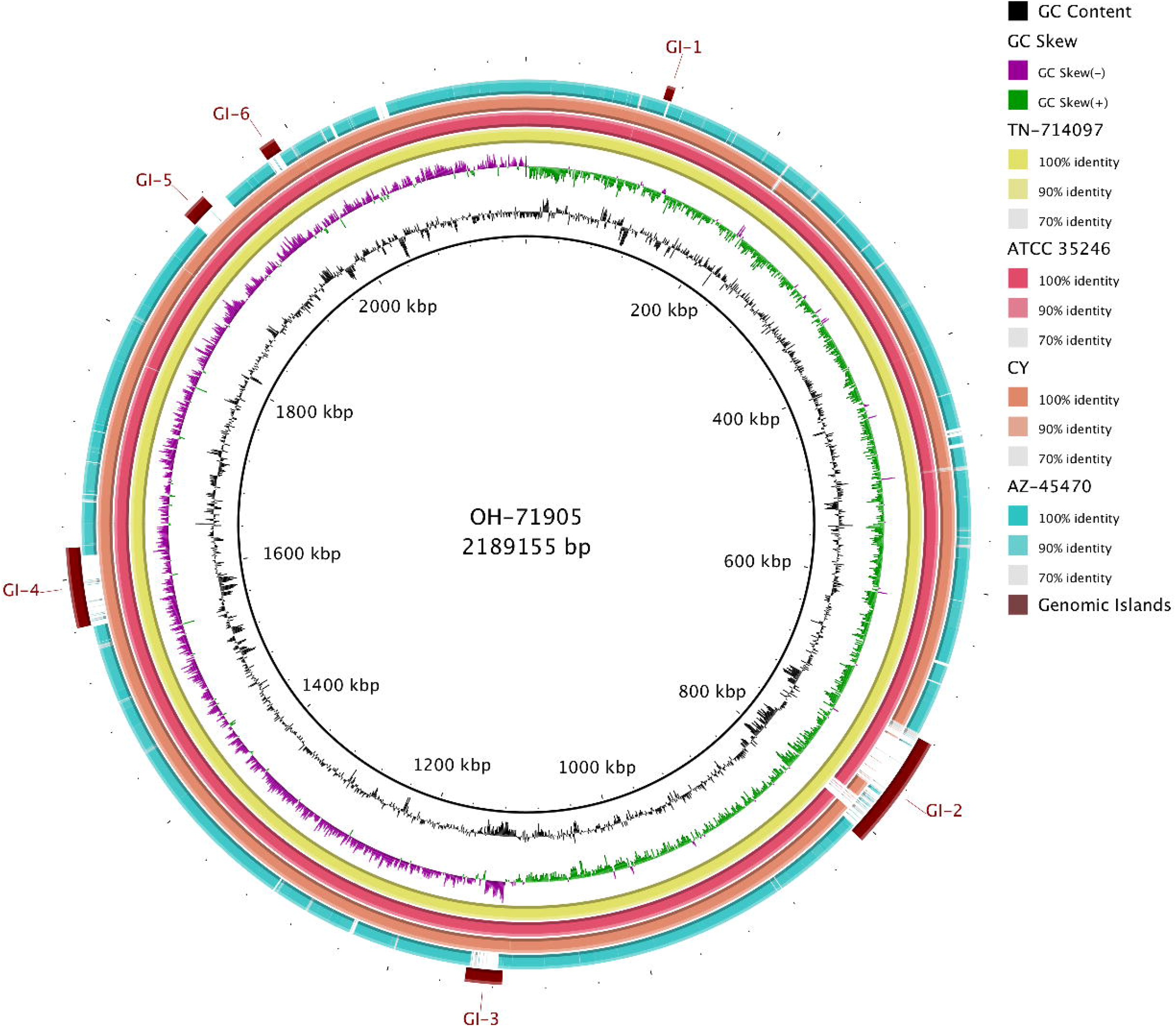
Comparative genome analysis between *S. zooepidemicus* isolates from pigs. Rings from outside to inside: 1) predicted genomic islands of OH-71905 (brown); 2) AZ-45470 from Arizona (blue); 3) CY from China (orange); 4) ATCC 35246 from China (Pink); 5) TN-714097 from Tennessee (yellow); 6) GC skew of OH-71905; 8) GC content of OH-71905; 9) OH-71905 genome. The blank spaces in the rings represented matches with less than 70% identity to the reference genome.

### Identification and distribution of genomic islands

A genomic island (GI) is part of a bacterial genome that has evidence of horizontal transmission origins. The GIs in OH-71905 and TN-74097 were predicted using IslandViewer 4. Six GIs with significantly different GC contents compared with the core genome were identified in both strains. The sizes of identified GIs varied greatly from 6 kb to 89 kb. Several GIs encode putative virulence genes and possibly contribute to the virulence of *S. zooepidemicus* (Appendix Table 4-5). GI-2 encodes a putative holing-like toxin and a putative type VI secretion system protein; GI-3 encodes a virulence associated protein E; GI-4 encodes a putative toxin PezT, a nucleotidyl transferase which belong to the AbiEii/AbiGii toxin family protein, and a putative VirB4 of type IV secretion system.

The distribution of the identified six GIs in all of the 45 *S. zooepidemicus* strains (Figure 4, Appendix Table 6) was assessed by examining the percentage of CDS present in six GIs. All six GIs were present (CDSs >75%, with more than 75% of the CDS in a GI present in other genomes) in all eight outbreak isolates from Ohio and Tennessee. Five complete GIs could be detected in ATCC 35246 and CY strains of the Chinese outbreaks, and only partial CDS (90/106, 84.91%) of GI-2 were found in ATCC 35246. In contrast, only one complete GI (GI-1) was present in swine isolate AZ-45470, which originated from a swine case that was not associated with high swine mortality outbreaks. More than 82% of CDS in all GIs except for GI-2 were present in the three isolates from human cases with guinea pig exposure (NVSLVA-S19, NVSLVA-S2, and NVSLVA-S22). In addition, the occurrence of five GIs (GI-2, GI-3, GI-4, GI-5, and GI-6) in other *S. zooepidemicus* isolates from human and other animal species was extremely low, two of 35 (5.71%) isolates possessed GI-3, GI-4 and GI-6 (CDS > 81%), while none of 35 isolates contained GI-2 or GI-5 (CDS < 61%).

**Figure 4.**
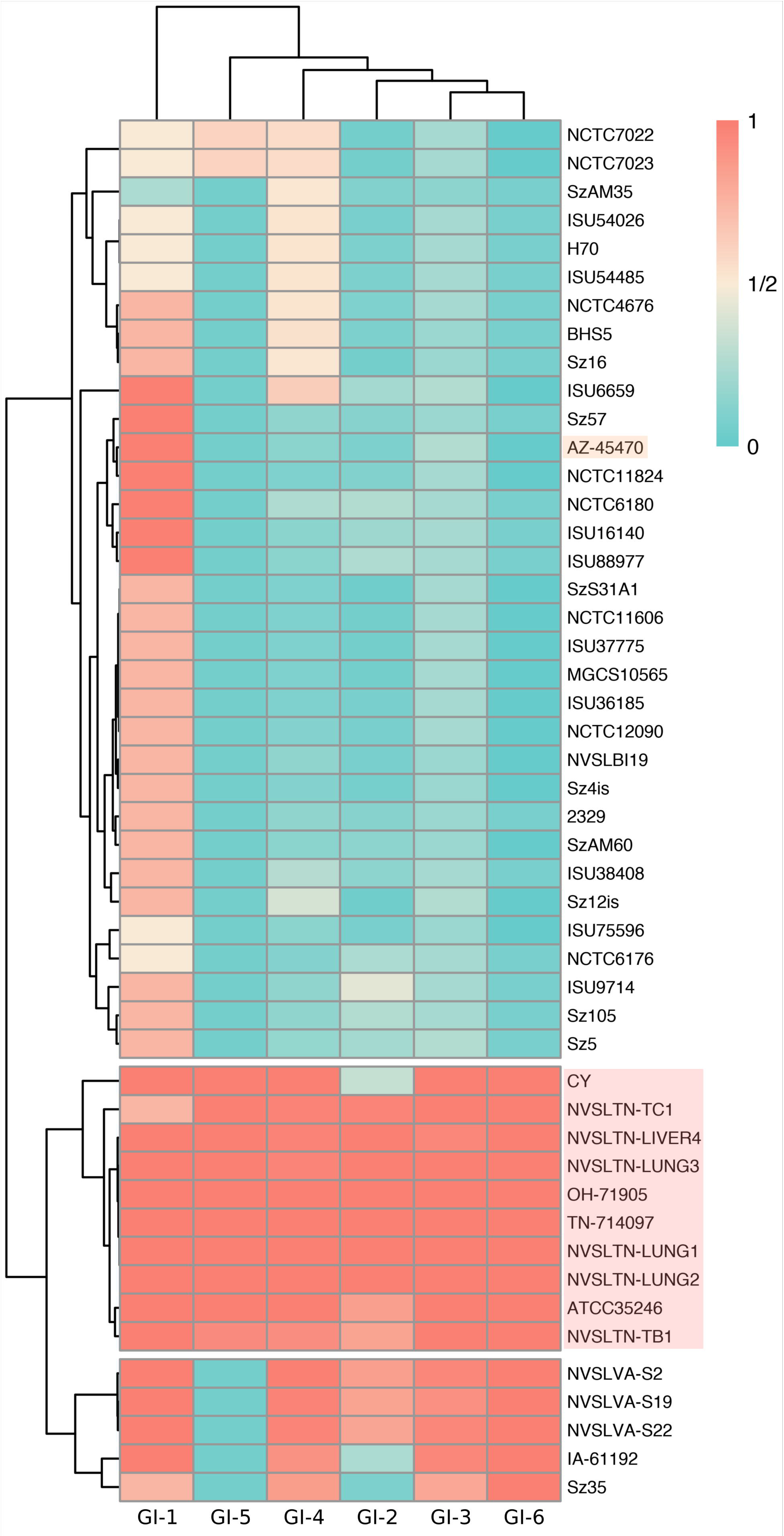
Two-way clustering of GIs prevalence among *S. zooepidemicus* isolates. A red-white-blue heat map was constructed based upon the percentage of the CDSs in each predicted OH-71905 GIs presented in all *S. zooepidemicus* isolates. Clustering was performed to illustrate the similarities between the prevalence of the GIs examined and between the *S. zooepidemicus* isolates with regard to CDS proportion. Red shade indicates porcine isolates associated with high mortality, while orange shade indicates another porcine isolate AZ-45470 without high mortality.

### Detection and distribution of putative virulence genes

The presence of 15 putative virulence genes, which have been previously reported in *S. zooepidemicus (13, 24-27),* was examined in all 48 isolates included in this study (Figure 5, Appendix Table 7). The M-like protein gene *szP* was present in all *S. zooepidemicus* isolates in our study albeit with variable sequences. The second M-like protein SzM and the newly identified Fic domain-containing protein BifA were recently reported virulence factors of *S. zooepidemicus*. Both genes were encoded by all eight isolates from Ohio and Tennessee outbreaks, the ATCC 35246 strain from China associated with high swine mortality and another swine strain from China (CY), but absent from the outbreak-unrelated swine isolate AZ-45470. The distribution of these two genes in other strains isolated from human or other animal species was only 18.42% (7/38). In addition, a fimbrial subunit protein coding gene *fszF* and a protective antigen-like protein coding gene *spaZ* were also present in all eight outbreak isolates from Ohio and Tennessee and the ATCC 35246 and CY strains from Chinese outbreaks, but negative in the outbreak-unrelated swine isolate AZ-45470. Several superantigen genes including *szeF, szeL, szeM, szeN* and *szeP* have been previously identified in *S. zooepidemicus* (*14*). These superantigen genes were all absent from all of the 11 swine isolates and infrequently present in other isolates from human and other animal species tested in our study (43.24%, 16/37).

**Figure 5.**
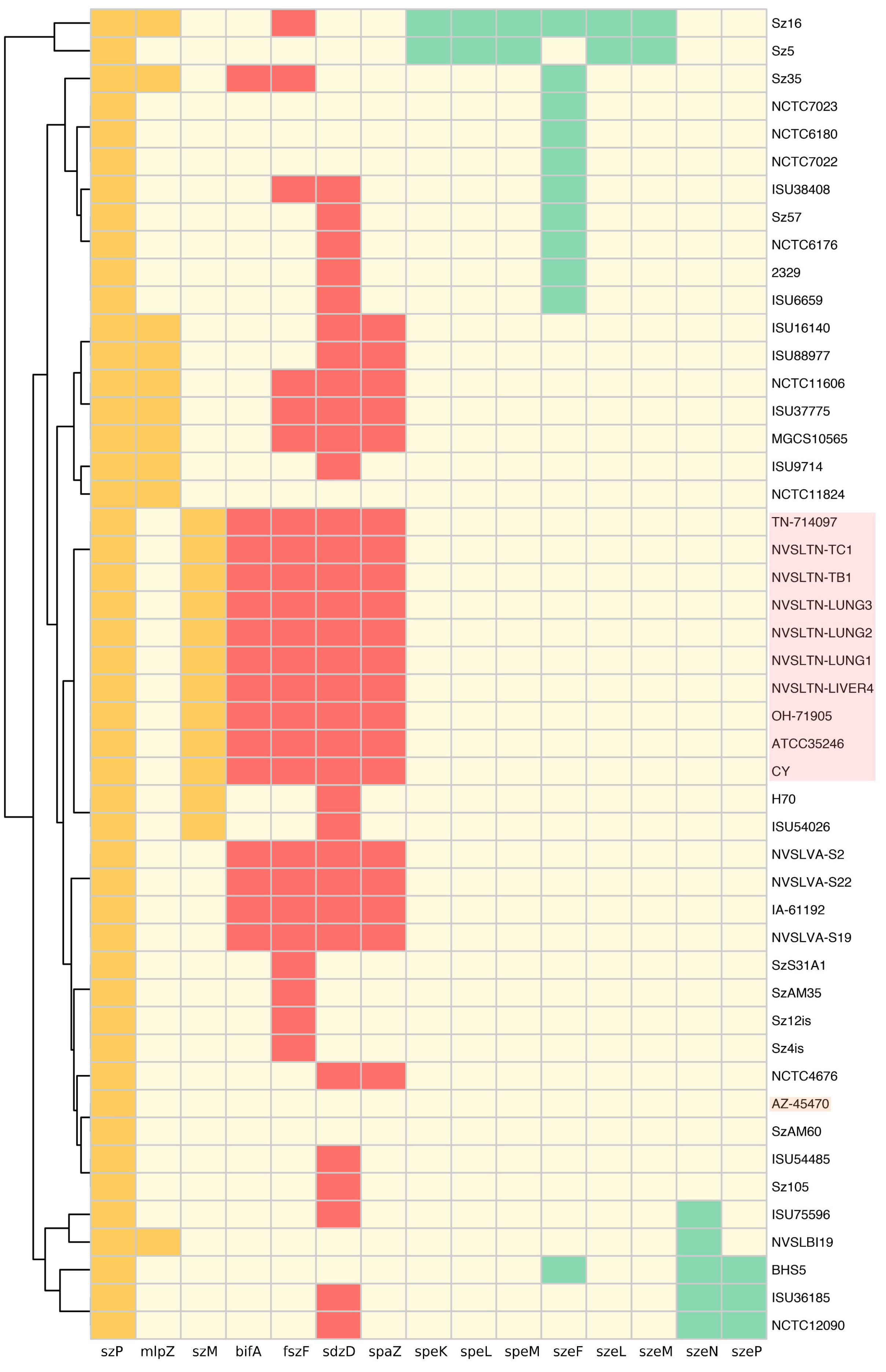
Distribution of putative virulence genes among 48 *S. zooepidemicus* isolates. Right side of the vertical line shows *S. zooepidemicus* isolates clustered by presence or absence of the putative virulence genes. Colored boxes indicate the presence of 3 classified groups: 1, M-like protein coding genes, yellow; 2, superantigen coding genes, green; 3, other putative virulence genes, red; white boxes indicate the absence of virulence genes. Red shade indicates porcine isolates associated with high mortality, while orange shade indicates another porcine isolate AZ-45470 without high mortality.

## Discussion

Although *S. zooepidemicus* is considered as an opportunistic pathogen in a large variety of host species including cats, rodents, mink, monkeys, and seals, the majority of cases were reported from domestic animals such as horses, dogs, ruminants, and pigs (*30*). *S. zooepidemicus* is a commensal organism in horses, but may act as an opportunistic pathogen causing abscesses, neonatal septicemia, and endometritis (*31*). Several studies revealed that *S. zooepidemicus* was the major causative agent of purulent infections in horses and foals leading to severe respiratory diseases (*32, 33). S. zooepidemicus* has also been reported as a causative agent of mastitis in ruminants inducing severe and deep tissue infections (*2*). In addition, emerging infections of *S. zooepidemicus* in dogs frequently occur as outbreaks causing severe and life-threatening diseases such as hemorrhagic pneumonia and septicemia (*34, 35*). In the swine industry, *S. zooepidemicus* has been a significant pathogen in some Asian countries with outbreaks reported in China (1975) where more than 300,000 pigs died (*15*) and in Indonesia nearly twenty years later (1994) (*4*). However, high mortality events due to *S. zooepidemicus* in swine have not previously been reported in the United States. Our genomic epidemiological analysis revealed that all eight *S. zooepidemicus* isolates from Ohio and Tennessee outbreaks were indistinguishable, yet clearly different from those isolated from horses, dogs, ruminants, and most other host species regardless of European or North American origin. These eight outbreak isolates were also significantly divergent from another outbreak-unrelated swine strain AZ-45470 from Arizona. What is especially concerning is that these eight outbreak isolates were clustered together with the ATCC 35246, the strain that caused the high mortality outbreak in China (*15*), and showed high similarity in their contents of genomic islands and virulence genes.

*S. zooepidemicus* is a known zoonotic pathogen and nearly 30 human cases of meningitis, septicemia, pneumonia, and glomerulonephritis have been documented (*36–41*). These human cases are usually associated with ingestion of animal products including unpasteurized milk and cheese or contact with companion animals such as horses, dogs, and guinea pigs (*30*). Transmission from pigs to human has, thus far, never been reported. Our results showed that the eight outbreak isolates of *S. zooepidemicus* in this study were classified as ST194, an ST type recorded in the *S. zooepidemicus* database from two human blood isolates recovered during 2001. Based on the whole genome phylogeny, the eight isolates were also closely clustered with three isolates from human patients who had severe clinical illness with guinea pig exposure (*6*), and moreover, their genomic islands and virulence genes were very similar. These results highlight significant public health concerns with these recent United States outbreak isolates. Swine producers, veterinarians, and other personnel who may directly or indirectly have contact with pigs should be aware of the potential of this organism to cause serious disease and death and *S. zooepidemicus* infection should be considered if they have purulent wounds or systemic symptoms of infection.

Our results also suggests the contributions of two previously identified virulence genes *szM* and *bifA* to the pathogenicity of *S. zooepidemicus*. Both genes were present in all eight *S. zooepidemicus* isolates and the ATCC 3546 strain, all of which have been involved in high mortalities. Conversely, both genes were absent from another outbreak-unrelated swine isolate AZ-45470, and were also extremely rare in strains from other animal species. It is possible that these two virulence genes are specifically associated with the hypervirulence of *S. zooepidemicus* to pigs, and further study of these specific genes is warranted. Superantigens are potent toxins, which may disrupt both innate and adaptive immune response and trigger non-specific T-cell proliferation and overzealous inflammatory production in the host (*42*). Several superantigen genes were shown to be significantly associated with non-strangles lymph node abscessation in the horse and probably contribute to virulence (*24*). However, in our study, all 11 swine isolates were negative for any superantigen gene, suggesting that they might not be necessary for the virulence of *S. zooepidemicus* in pigs. In addition, many deletions were identified from the genome of the outbreak-unrelated swine isolate AZ-45470 compared with those of eight *S. zooepidemicus* outbreak isolates and the ATCC 3546 strain. Among them, six large deletions were predicted genomic islands that were absent from this strain and several of these islands encoded putative virulence genes including putative toxin genes and type IV and VI secretion system proteins. It will be very interesting to determine if these genes and islands would contribute to the virulence of *S. zooepidemicus* in pigs.

In summary, we performed genomic epidemiological and comparative genomic analyses with *S. zooepidemicus* isolates associated with high swine mortality in the United States. Our findings provide significant and timely insights for a better understanding of the epidemiology and virulence of *S. zooepidemicus* isolates associated with highly unexpected and severe outbreaks that occurred very recently in the US swine population. In addition, identification of specific virulence genes and genomic islands may lead to the development of novel molecular diagnostic tools, and provide the basis for future investigation of virulence mechanisms and control measures.

Ms. Chen is a graduate student in the College of Veterinary Medicine at Iowa State University. Her Primary research interests include molecular epidemiology and bacterial pathogensis.

## Acknowledgement

This study was partially supported by the Swine Health Information Center (SHIC#19-236). We thank Ying Zheng and Dr. Huigang Shen for excellent technical assistance.

