## Supplemental tables for "Genetic characterization of *Streptococcus equi* subspecies *zooepidemicus* associated with high swine mortality in United States"

**Appendix**

**Appendix Table 1**. Basic information for 24 *S. zooepidemicus* isolates for whole genome sequencing.

| Isolates | Host | Date | Country | SRA accession | Contigs and genome accession | Assembly level |
| --- | --- | --- | --- | --- | --- | --- |
| ISU37775 | caprine | 2016 | USA: Iowa | SRR10579788 | WMAE00000000 | Contig |
| ISU6659 | chinchilla | 2017 | USA: Iowa | SRR10579787 | WMAH00000000 | Contig |
| ISU54485 | feline | 2017 | USA: Illinois | SRR10579782 | WMAB00000000 | Contig |
| ISU9714 | canine | 2018 | USA: Iowa | SRR10579781 | WNGY00000000 | Contig |
| ISU36185 | equine | 2018 | USA: Iowa | SRR10579780 | WMAF00000000 | Contig |
| ISU38408 | equine | 2018 | USA: Iowa | SRR10579779 | WMAD00000000 | Contig |
| ISU75596 | feline | 2018 | USA: Iowa | SRR10579778 | WMAA00000000 | Contig |
| ISU88977 | feline | 2018 | USA: Iowa | SRR10579777 | WLZZ00000000 | Contig |
| ISU16140 | equine | 2019 | USA: Iowa | SRR10579776 | WMAG00000000 | Contig |
| ISU54026 | equine | 2019 | USA: Iowa | SRR10579786 | WMAC00000000 | Contig |
| AZ-45470 | porcine | 2019 | USA: Arizona | SRR10579775 | CP046041 | Complete Genome |
| IA-61192 | equine | 2019 | USA: Iowa | SRR10579785 | WOFY00000000 | Contig |
| OH-71905 | porcine | 2019 | USA: Ohio | SRR10579784 | CP046040 | Complete Genome |
| TN-74097 | porcine | 2019 | USA: Tennessee | SRR10579783 | CP046042 | Complete Genome |
| NVSLBI19 | equine | 2019 | USA: Tennessee | SRX7265086 | WMAI00000000 | Contig |
| NVSLTN-LIVER4 | porcine | 2019 | USA: Tennessee | SRX7265087 | WOFZ00000000 | Contig |
| NVSLTN-LUNG1 | porcine | 2019 | USA: Tennessee | SRX7265088 | WOGA00000000 | Contig |
| NVSLTN-LUNG2 | porcine | 2019 | USA: Tennessee | SRX7265089 | WOGB00000000 | Contig |
| NVSLTN-LUNG3 | porcine | 2019 | USA: Tennessee | SRX7265090 | WOGC00000000 | Contig |
| NVSLTN-TB1 | porcine | 2019 | USA: Tennessee | SRX7265091 | WOGD00000000 | Contig |
| NVSLTN-TC1 | porcine | 2019 | USA: Tennessee | SRX7265092 | WOGE00000000 | Contig |
| NVSLVA-S2 | guinea pig | 2013 | USA: Virginia | SRX7200846 | WOGF00000000 | Contig |
| NVSLVA-S19 | guinea pig | 2013 | USA: Virginia | SRX7200845 | WOGG00000000 | Contig |
| NVSLVA-S22 | guinea pig | 2013 | USA: Virginia | SRX7200844 | WOGH00000000 | Contig |

**Appendix Table 2**. Basic information for 24 *S. zooepidemicus* isolates downloaded from NCBI

| Isolates | Host | Date | Country | Contigs and genome accession | Assembly level |
| --- | --- | --- | --- | --- | --- |
| 2329 | equine | 1996 | United Kingdom | GCA_000836615.1 | Contig |
| ATCC 35246 | porcine | 1975 | China | CP002904.1 | Complete Genome |
| BHS5 | canine | 1999 | United Kingdom | GCA_000208725.1 | Scaffold |
| CY | porcine | 1998 | China: Nanjing | [CP006770.1](https://www.ncbi.nlm.nih.gov/nuccore/CP006770.1) | Complete Genome |
| H70 | equine | 2000 | United Kingdom: Newmarket | [FM204884.1](https://www.ncbi.nlm.nih.gov/nuccore/FM204884.1) | Complete Genome |
| MGCS10565 |  | 2008 | Brazil | [CP001129.1](https://www.ncbi.nlm.nih.gov/nuccore/CP001129.1) | Complete Genome |
| NCTC4676 | cattle | 1900/1936 | United Kingdom: London | GCA_900459475.1 | Contig |
| NCTC6176 | mouse | 1900/1941 | United Kingdom | [LS483368.1](https://www.ncbi.nlm.nih.gov/nuccore/LS483368.1) | Complete Genome |
| NCTC6180 | equine | 1900/1941 |  | [LR134317.1](https://www.ncbi.nlm.nih.gov/nuccore/LR134317.1) | Complete Genome |
| NCTC7022 |  | 1945 | United Kingdom: Weybridge | [LS483325.1](https://www.ncbi.nlm.nih.gov/nuccore/LS483325.1) | Complete Genome |
| NCTC7023 |  | 1945 | United Kingdom | GCA_900460185.1 | Contig |
| NCTC11606 |  | 1983 | United Kingdom: North Yorkshire | [LS483354.1](https://www.ncbi.nlm.nih.gov/nuccore/LS483354.1) | Complete Genome |
| NCTC11824 | human | 1980 | United Kingdom: Blackburn | [LS483380.1](https://www.ncbi.nlm.nih.gov/nuccore/LS483380.1) | Complete Genome |
| NCTC12090 |  | 1987 | United Kingdom: Northallerton | [LS483328.1](https://www.ncbi.nlm.nih.gov/nuccore/LS483328.1) | Complete Genome |
| Sz4is | equine | 2007 | Iceland | GCA_000876215.1 | Contig |
| Sz5 | equine | 2007 | Denmark: Ansager | GCA_000876355.1 | Contig |
| Sz12is | equine | 2007 | Iceland | GCA_000876305.1 | Contig |
| Sz16 | equine | 2007 | Denmark: Ansager | GCA_000876295.1 | Contig |
| Sz35 | equine | 2007 | Denmark: Ansager | GCA_000876365.1 | Contig |
| Sz57 | equine | 2007 | Denmark: Ansager | GCA_000876375.1 | Contig |
| Sz105 | equine | 2007 | Denmark: Ansager | GCA_000876195.1 | Contig |
| SzAM35 | equine | 2007 | USA: Lexington, Kentucky | GCA_000876275.1 | Contig |
| SzAM60 | equine | 2007 | USA: Lexington, Kentucky | GCA_000876285.1 | Contig |
| SzS31A1 | equine |  | Denmark | GCA_000445225.2 | Contig |

**Appendix Table 3**. Detailed description of MLST analysis for 48 *S. zooepidemicus* isolates

| Isolates | MLST typing | | | | | | | |
| --- | --- | --- | --- | --- | --- | --- | --- | --- |
|  | arcC | nrdE | proS | spi | tdk | tpi | yqiL | ST type |
| ISU37775 | 3 | no matchs | 5 | 6 | 7 | 7 | 9 | unknown |
| ISU6659 | 13 | 4 | 11 | 25 | 1 | 13 | 10 | unknown |
| ISU54485 | 3 | 3 | 1 | 6 | 1 | 5 | 16 | 40 |
| ISU9714 | 3 | 3 | 15 | 7 | 10 | 18 | 30 | 384 |
| ISU36185 | 5 | 3 | 3 | 5 | 3 | no matchs | 16 | unknown |
| ISU38408 | 2 | 3 | 2 | 14 | 1 | 22 | no matchs | unknown |
| ISU75596 | 5 | 3 | 5 | 5 | 3 | 14 | 16 | 70 |
| ISU88977 | 9 | 3 | 38 | 40 | no matchs | 14 | 16 | unknown |
| ISU16140 | 9 | 3 | 38 | 40 | 1 | 14 | 16 | 222 |
| ISU54026 | 10 | 14 | 22 | 20 | 1 | 10 | 6 | 340 |
| AZ-45470 | 1 | 19 | 1 | 1 | 1 | 1 | 1 | 71 |
| IA-61192 | 1 | 3 | 36 | 45 | 1 | 15 | 29 | 329 |
| OH-71905 | 27 | 3 | 1 | 45 | 1 | 34 | 46 | 194 |
| TN-74097 | 27 | 3 | 1 | 45 | 1 | 34 | 46 | 194 |
| NVSLBI19 | 6 | 6 | 6 | 4 | 5 | 7 | 7 | 8 |
| NVSLTN-LIVER4 | 27 | 3 | 1 | 45 | 1 | 34 | 46 | 194 |
| NVSLTN-LUNG1 | 27 | 3 | 1 | 45 | 1 | 34 | 46 | 194 |
| NVSLTN-LUNG2 | 27 | 3 | 1 | 45 | 1 | 34 | 46 | 194 |
| NVSLTN-LUNG3 | 27 | 3 | 1 | 45 | 1 | 34 | 46 | 194 |
| NVSLTN-TB1 | 27 | 3 | 1 | 45 | 1 | 34 | 46 | 194 |
| NVSLTN-TC1 | 27 | 3 | 1 | 45 | 1 | 34 | 46 | 194 |
| NVSLVA-S2 | 27 | 19 | 15 | 45 | 1 | 34 | 46 | unknown |
| NVSLVA-S19 | 27 | 19 | 15 | 45 | 1 | 34 | 46 | unknown |
| NVSLVA-S22 | 27 | 19 | 15 | 45 | 1 | 34 | 46 | unknown |
| 2329 | 9 |  | 9 | 4 | 1 | 11 | 1 | unknown |
| ATCC 35246 | 27 | 3 | 1 | 45 | 1 | 34 | 46 | 194 |
| BHS5 | 21 | 3 | 12 | 29 | 1 | 45 | 15 | 123 |
| CY | 27 | 3 | 1 | 45 | 1 | 34 | 46 | 194 |
| H70 | 1 | 1 | 1 | 1 | 1 | 1 | 1 | 1 |
| MGCS10565 | 3 | 10 | 5 | 6 | 7 | 7 | 9 | 72 |
| NCTC4676 | 3 | 3 | 4 | 45 | 1 | 5 | 16 | 214 |
| NCTC6176 | 3 | 2 | 5 | 19 | 14 | 5 | 28 | 391 |
| NCTC6180 | 9 | 18 | 64 | 6 | 1 | 47 | 70 | unknown |
| NCTC7022 | 1 | 11 | 20 | 4 | 24 | 16 | 3 | 219 |
| NCTC7023 | 1 | 11 | 20 | 4 | 24 | 16 | 3 | 219 |
| NCTC11606 | 3 | 10 | 5 | 6 | 7 | 7 | 9 | 72 |
| NCTC11824 | 10 | 14 | 22 | 14 | 1 | 10 | 15 | 306 |
| NCTC12090 | 5 | 3 | 3 | 5 | 3 | 5 | 5 | 5 |
| Sz4is | 2 | 11 | 16 | 26 | 7 | 9 | 6 | 279 |
| Sz5 | 18 | 22 | 54 | 45 | 1 | 7 | 2 | 303 |
| Sz12is | 2 | 11 | 16 | 26 | 7 | 9 | 6 | 279 |
| Sz16 | 4 | 3 | 6 | 2 | 1 | 19 | 26 | 156 |
| Sz35 | 2 | 4 | 4 | 7 | 4 | 5 | 43 | 203 |
| Sz57 | 2 | 6 | 14 | 4 | 4 | 14 | 16 | 96 |
| Sz105 | 3 | 3 | 10 | 22 | 10 | 5 | 12 | 140 |
| SzAM35 | 4 | 3 | 5 | 2 | 1 | 19 | 26 | 65 |
| SzAM60 | no matchs | 11 | 15 | 6 | 14 | 16 | 16 | unknown |
| SzS31A1 | 2 | 11 | 16 | 26 | 7 | 9 | 6 | 279 |

**Appendix Table 4.** General feature for 6 predicted genomic islands of OH-71905

| Island | Island start | Island end | Length |
| --- | --- | --- | --- |
| GI-1 | 108516 | 115047 | 6531 |
| GI-2 | 720854 | 810293 | 89439 |
| GI-3 | 1113013 | 1141475 | 28462 |
| GI-4 | 1562548 | 1624048 | 61500 |
| GI-5 | 1897996 | 1918020 | 20024 |
| GI-6 | 1973842 | 1987041 | 13199 |

**Appendix Table 5**. Detailed description for CDSs in predicted genomic islands of OH-71905

| Gene name | Gene ID | Locus | Gene start | Gene end | Strand | Product |
| --- | --- | --- | --- | --- | --- | --- |
| GI-1 | | | | | | |
|  |  | GJS32_00640 | 108516 | 108726 | -1 | ISL3 family transposase |
| QGM13048.1 |  | GJS32_00645 | 108996 | 109523 | 1 | hypothetical protein |
| QGM13049.1 |  | GJS32_00650 | 110075 | 111133 | 1 | alanine racemase |
| QGM13050.1 |  | GJS32_00655 | 111194 | 111559 | 1 | DUF1033 family protein |
| QGM13051.1 |  | GJS32_00660 | 111806 | 112678 | 1 | helix-turn-helix domain-containing protein |
| QGM13052.1 |  | GJS32_00665 | 113083 | 115047 | 1 | kinase |
| GI-2 | | | | | | |
| QGM13553.1 |  | GJS32_03415 | 720854 | 722215 | 1 | bacteriocin secretion accessory protein |
| QGM13554.1 |  | GJS32_03420 | 722594 | 722866 | 1 | hypothetical protein |
| QGM13555.1 |  | GJS32_03425 | 722866 | 723531 | 1 | CPBP family intramembrane metalloprotease |
| QGM13556.1 |  | GJS32_03430 | 723809 | 723973 | 1 | hypothetical protein |
| QGM13557.1 |  | GJS32_03435 | 724866 | 727175 | 1 | DUF4135 domain-containing protein |
| QGM13558.1 |  | GJS32_03440 | 727165 | 727878 | 1 | hypothetical protein |
| QGM13559.1 |  | GJS32_03445 | 727893 | 728069 | 1 | hypothetical protein |
| QGM13560.1 |  | GJS32_03450 | 728085 | 728279 | 1 | hypothetical protein |
| QGM13561.1 |  | GJS32_03455 | 728295 | 728477 | 1 | hypothetical protein |
| QGM13562.1 | tnpA | GJS32_03460 | 728700 | 729173 | 1 | IS200/IS605 family transposase |
| QGM13563.1 | tnpA | GJS32_03465 | 729343 | 729807 | 1 | IS200/IS605 family transposase |
| QGM13564.1 |  | GJS32_03470 | 729951 | 730172 | -1 | helix-turn-helix domain-containing protein |
| QGM13565.1 |  | GJS32_03475 | 730162 | 731820 | -1 | DUF2326 domain-containing protein |
| QGM13566.1 |  | GJS32_03480 | 731811 | 732035 | -1 | hypothetical protein |
| QGM13567.1 |  | GJS32_03485 | 732035 | 733153 | -1 | hypothetical protein |
| QGM13568.1 |  | GJS32_03490 | 733586 | 734182 | 1 | hypothetical protein |
| QGM13569.1 |  | GJS32_03495 | 734390 | 734899 | 1 | type III toxin-antitoxin system ToxN/AbiQ family toxin |
| QGM13570.1 |  | GJS32_03500 | 734944 | 736899 | -1 | hypothetical protein |
| QGM13571.1 |  | GJS32_03505 | 737134 | 737697 | -1 | DUF2815 family protein |
| QGM13572.1 |  | GJS32_03510 | 737702 | 738823 | -1 | DUF2800 domain-containing protein |
| QGM13573.1 |  | GJS32_03515 | 738816 | 739139 | -1 | hypothetical protein |
| QGM13574.1 |  | GJS32_03520 | 739500 | 741788 | 1 | DNA primase |
| QGM13575.1 |  | GJS32_03525 | 742069 | 742350 | 1 | VRR-NUC domain-containing protein |
| QGM13576.1 |  | GJS32_03530 | 742331 | 743707 | 1 | DEAD/DEAH box helicase |
| QGM13577.1 |  | GJS32_03535 | 743700 | 744176 | 1 | hypothetical protein |
| QGM13578.1 |  | GJS32_03540 | 744371 | 744553 | 1 | hypothetical protein |
| QGM13579.1 |  | GJS32_03545 | 744601 | 745638 | 1 | methionine adenosyltransferase |
| QGM13580.1 |  | GJS32_03550 | 745640 | 745999 | 1 | HNH endonuclease |
| QGM13581.1 |  | GJS32_03555 | 746039 | 746584 | 1 | hypothetical protein |
| QGM13582.1 |  | GJS32_03560 | 746764 | 747219 | 1 | helix-turn-helix domain-containing protein |
| QGM13583.1 |  | GJS32_03565 | 747191 | 748438 | 1 | DNA modification methylase |
| QGM13584.1 |  | GJS32_03570 | 748443 | 748916 | 1 | phage terminase small subunit P27 family |
| QGM13585.1 |  | GJS32_03575 | 748913 | 750505 | 1 | terminase large subunit |
| QGM13586.1 |  | GJS32_03580 | 750576 | 750833 | 1 | type II toxin-antitoxin system Phd/YefM family antitoxin |
| QGM13587.1 |  | GJS32_03585 | 750830 | 751201 | 1 | type II toxin-antitoxin system death-on-curing family toxin |
|  |  | GJS32_03590 | 751624 | 751813 | 1 | hypothetical protein |
| QGM13588.1 |  | GJS32_03595 | 751786 | 751974 | 1 | hypothetical protein |
| QGM13589.1 |  | GJS32_03600 | 752031 | 753320 | 1 | phage portal protein |
| QGM13590.1 |  | GJS32_03605 | 753313 | 754011 | 1 | Clp protease ClpP |
| QGM13591.1 |  | GJS32_03610 | 754025 | 755230 | 1 | phage major capsid protein |
| QGM13592.1 |  | GJS32_03615 | 755230 | 755487 | 1 | phage gp6-like head-tail connector protein |
| QGM13593.1 |  | GJS32_03620 | 755487 | 755825 | 1 | head-tail adaptor protein |
| QGM13594.1 |  | GJS32_03625 | 755818 | 756186 | 1 | HK97 gp10 family phage protein |
| QGM13595.1 |  | GJS32_03630 | 756195 | 756521 | 1 | hypothetical protein |
| QGM13596.1 |  | GJS32_03635 | 756524 | 757093 | 1 | phage tail protein |
| QGM13597.1 |  | GJS32_03640 | 757105 | 757524 | 1 | hypothetical protein |
| QGM13598.1 |  | GJS32_03645 | 757707 | 760826 | 1 | phage tail tape measure protein |
| QGM13599.1 |  | GJS32_03650 | 760823 | 761548 | 1 | phage tail protein |
| QGM14805.1 |  | GJS32_03655 | 761548 | 764463 | 1 | carbamoylsarcosine amidase |
| QGM13600.1 |  | GJS32_03660 | 764476 | 766332 | 1 | hypothetical protein |
| QGM13601.1 |  | GJS32_03665 | 766350 | 766754 | 1 | holin |
| QGM13602.1 |  | GJS32_03670 | 766756 | 768225 | 1 | LysM peptidoglycan-binding domain-containing protein |
| QGM13603.1 |  | GJS32_03675 | 768380 | 768487 | 1 | putative holin-like toxin |
| QGM13604.1 |  | GJS32_03680 | 768864 | 770105 | 1 | recombinase family protein |
| QGM14806.1 |  | GJS32_03685 | 770044 | 771360 | 1 | recombinase family protein |
| QGM13605.1 |  | GJS32_03690 | 771464 | 773086 | 1 | DUF2075 domain-containing protein |
| QGM13606.1 |  | GJS32_03695 | 773187 | 773375 | 1 | hypothetical protein |
| QGM14807.1 |  | GJS32_03700 | 773382 | 773663 | 1 | hypothetical protein |
|  |  | GJS32_03705 | 773678 | 774029 | 1 | replication initiation protein |
|  |  | GJS32_03710 | 774023 | 774531 | 1 | nucleoside triphosphate hydrolase |
| QGM13607.1 |  | GJS32_03715 | 774528 | 775001 | 1 | DUF3801 domain-containing protein |
| QGM13608.1 |  | GJS32_03720 | 774998 | 775336 | 1 | conjugal transfer protein |
| QGM13609.1 |  | GJS32_03725 | 775387 | 775524 | 1 | hypothetical protein |
| QGM13610.1 |  | GJS32_03730 | 775524 | 776048 | 1 | resolvase |
|  |  | GJS32_03735 | 776045 | 777166 | 1 | DUF4368 domain-containing protein |
| QGM14808.1 |  | GJS32_03740 | 777350 | 777454 | 1 | hypothetical protein |
| QGM13611.1 |  | GJS32_03745 | 777526 | 777846 | 1 | DUF3847 domain-containing protein |
| QGM13612.1 |  | GJS32_03750 | 778086 | 779543 | 1 | conjugal transfer protein TraA |
| QGM13613.1 |  | GJS32_03755 | 779650 | 780183 | 1 | DUF1697 domain-containing protein |
|  |  | GJS32_03760 | 780326 | 780577 | 1 | type IV secretory system conjugative DNA transfer family protein |
| QGM13614.1 |  | GJS32_03765 | 780620 | 782290 | 1 | DUF4368 domain-containing protein |
| QGM13615.1 |  | GJS32_03770 | 782350 | 782637 | 1 | 50S ribosomal protein L7/L12 |
| QGM13616.1 |  | GJS32_03775 | 782764 | 783084 | 1 | DUF3847 domain-containing protein |
| QGM13617.1 |  | GJS32_03780 | 783322 | 784806 | 1 | conjugal transfer protein TraA |
| QGM13618.1 |  | GJS32_03785 | 784925 | 785341 | 1 | hypothetical protein |
| QGM13619.1 |  | GJS32_03790 | 785344 | 785547 | 1 | helix-turn-helix domain-containing protein |
| QGM13620.1 |  | GJS32_03795 | 785651 | 785815 | 1 | hypothetical protein |
| QGM13621.1 |  | GJS32_03800 | 785890 | 786351 | 1 | conjugal transfer protein TraG |
| QGM13622.1 |  | GJS32_03805 | 786383 | 786583 | 1 | hypothetical protein |
| QGM13623.1 |  | GJS32_03810 | 786608 | 788458 | 1 | DUF4368 domain-containing protein |
| QGM13624.1 |  | GJS32_03815 | 788531 | 789055 | 1 | PadR family transcriptional regulator |
| QGM13625.1 |  | GJS32_03820 | 789059 | 789709 | 1 | HEAT repeat domain-containing protein |
| QGM13626.1 |  | GJS32_03825 | 789693 | 790157 | 1 | hypothetical protein |
| QGM13627.1 |  | GJS32_03830 | 790337 | 790753 | 1 | sigma-70 family RNA polymerase sigma factor |
| QGM13628.1 |  | GJS32_03835 | 790822 | 791088 | 1 | DUF3847 domain-containing protein |
| QGM14809.1 |  | GJS32_03840 | 791634 | 792803 | -1 | IS110 family transposase |
| QGM13629.1 |  | GJS32_03845 | 793063 | 794700 | 1 | MobA/MobL protein |
|  |  | GJS32_03850 | 794908 | 795522 | 1 | replisome organizer |
|  |  | GJS32_03855 | 795513 | 795743 | 1 | transposase |
| QGM13630.1 |  | GJS32_03860 | 795926 | 796783 | 1 | PrsW family intramembrane metalloprotease |
| QGM13631.1 |  | GJS32_03865 | 797114 | 797374 | 1 | hypothetical protein |
| QGM13632.1 |  | GJS32_03870 | 797695 | 798348 | 1 | hypothetical protein |
| QGM13633.1 |  | GJS32_03875 | 798577 | 798762 | 1 | hypothetical protein |
|  |  | GJS32_03880 | 798774 | 799127 | 1 | type VI secretion protein |
| QGM13634.1 | ltrA | GJS32_03885 | 799879 | 801789 | 1 | group II intron reverse transcriptase/maturase |
| QGM14810.1 |  | GJS32_03890 | 801812 | 802402 | 1 | TraM recognition domain-containing protein |
| QGM13635.1 |  | GJS32_03895 | 802483 | 803031 | 1 | hypothetical protein |
| QGM13636.1 |  | GJS32_03900 | 803078 | 803296 | 1 | hypothetical protein |
|  |  | GJS32_03905 | 803356 | 803997 | 1 | hypothetical protein |
| QGM13637.1 |  | GJS32_03910 | 804136 | 804510 | 1 | hypothetical protein |
| QGM13638.1 |  | GJS32_03915 | 804687 | 805892 | 1 | AAA family ATPase |
| QGM13639.1 |  | GJS32_03920 | 806728 | 807372 | -1 | histidine phosphatase family protein |
| QGM13640.1 |  | GJS32_03925 | 807391 | 807756 | -1 | YccF domain-containing protein |
| QGM13641.1 |  | GJS32_03930 | 807753 | 808232 | -1 | aminoacyl-tRNA deacylase |
| QGM13642.1 | thiT | GJS32_03935 | 808393 | 808956 | -1 | energy-coupled thiamine transporter ThiT |
| QGM13643.1 |  | GJS32_03940 | 809427 | 810293 | -1 | N-acetylmuramoyl-L-alanine amidase |
| GI-3 | | | | | | |
| QGM13895.1 |  | GJS32_05345 | 1113013 | 1113450 | -1 | GtrA family protein |
|  |  | GJS32_05350 | 1113558 | 1114503 | -1 | IS30 family transposase |
| QGM13896.1 |  | GJS32_05355 | 1114722 | 1115420 | 1 | helix-turn-helix domain-containing protein |
| QGM13897.1 |  | GJS32_05360 | 1115751 | 1116545 | -1 | hypothetical protein |
| QGM13898.1 |  | GJS32_05365 | 1116909 | 1117787 | -1 | DUF4868 domain-containing protein |
| QGM13899.1 |  | GJS32_05370 | 1117762 | 1118235 | -1 | hypothetical protein |
|  |  | GJS32_05375 | 1118300 | 1121462 | -1 | HsdR family type I site-specific deoxyribonuclease |
| QGM14830.1 |  | GJS32_05380 | 1121471 | 1122373 | -1 | hypothetical protein |
| QGM13900.1 |  | GJS32_05385 | 1122392 | 1123324 | -1 | restriction endonuclease subunit S |
|  |  | GJS32_05390 | 1123498 | 1125086 | -1 | N-6 DNA methylase |
| QGM13901.1 |  | GJS32_05395 | 1125103 | 1125309 | -1 | helix-turn-helix domain-containing protein |
| QGM13902.1 |  | GJS32_05400 | 1125468 | 1126592 | -1 | relaxase/mobilization nuclease domain-containing protein |
| QGM13903.1 |  | GJS32_05405 | 1126711 | 1128513 | -1 | group II intron reverse transcriptase/maturase |
|  |  | GJS32_05410 | 1129086 | 1129412 | -1 | relaxase/mobilization nuclease domain-containing protein |
| QGM13904.1 | mobC | GJS32_05415 | 1129412 | 1129792 | -1 | plasmid mobilization relaxosome protein MobC |
| QGM13905.1 |  | GJS32_05420 | 1130113 | 1131504 | -1 | virulence-associated protein E |
| QGM13906.1 |  | GJS32_05425 | 1131497 | 1132426 | -1 | DNA primase |
| QGM13907.1 |  | GJS32_05430 | 1132534 | 1132917 | -1 | hypothetical protein |
| QGM14831.1 |  | GJS32_05435 | 1132910 | 1133986 | -1 | tyrosine-type recombinase/integrase |
| QGM13908.1 |  | GJS32_05440 | 1133993 | 1134172 | -1 | hypothetical protein |
| QGM13909.1 |  | GJS32_05445 | 1134281 | 1135540 | -1 | ISL3-like element ISSeq1 family transposase |
| QGM13910.1 |  | GJS32_05450 | 1135746 | 1136918 | 1 | hypothetical protein |
|  |  | GJS32_05455 | 1136932 | 1137872 | 1 | phosphoribosylaminoimidazolesuccinocarboxamide synthase |
| QGM13911.1 |  | GJS32_05460 | 1137884 | 1138762 | 1 | hypothetical protein |
| QGM13912.1 | guaA | GJS32_05465 | 1138928 | 1140490 | -1 | glutamine-hydrolyzing GMP synthase |
| QGM13913.1 |  | GJS32_05470 | 1140777 | 1141475 | 1 | UTRA domain-containing protein |
| GI-4 | | | | | | |
|  |  | GJS32_07405 | 1561119 | 1562555 | -1 | recombinase |
| QGM14264.1 |  | GJS32_07410 | 1562548 | 1564161 | -1 | recombinase |
|  |  | GJS32_07415 | 1564161 | 1565468 | -1 | recombinase |
| QGM14265.1 |  | GJS32_07420 | 1565592 | 1566851 | 1 | ISL3-like element ISSeq1 family transposase |
| QGM14266.1 |  | GJS32_07425 | 1566860 | 1567042 | -1 | hypothetical protein |
| QGM14267.1 |  | GJS32_07430 | 1567061 | 1567252 | -1 | hypothetical protein |
| QGM14268.1 |  | GJS32_07435 | 1567382 | 1570027 | -1 | AAA family ATPase |
| QGM14269.1 |  | GJS32_07440 | 1570032 | 1570718 | -1 | hypothetical protein |
| QGM14270.1 |  | GJS32_07445 | 1570711 | 1571982 | -1 | hypothetical protein |
| QGM14271.1 |  | GJS32_07450 | 1571985 | 1572755 | -1 | toxin PezT |
| QGM14272.1 |  | GJS32_07455 | 1572755 | 1573231 | -1 | helix-turn-helix domain-containing protein |
| QGM14273.1 |  | GJS32_07460 | 1573301 | 1573588 | -1 | hypothetical protein |
| QGM14274.1 |  | GJS32_07465 | 1573633 | 1574028 | -1 | chemotaxis protein |
| QGM14275.1 |  | GJS32_07470 | 1574032 | 1574259 | -1 | hypothetical protein |
|  |  | GJS32_07475 | 1574313 | 1575399 | -1 | DNA primase |
| QGM14854.1 |  | GJS32_07480 | 1575438 | 1576076 | -1 | peptidylprolyl isomerase |
| QGM14276.1 |  | GJS32_07485 | 1576259 | 1577773 | -1 | ATP-binding cassette domain-containing protein |
| QGM14277.1 |  | GJS32_07490 | 1577757 | 1578776 | -1 | radical SAM protein |
| QGM14278.1 |  | GJS32_07495 | 1579065 | 1580114 | -1 | radical SAM protein |
| QGM14279.1 |  | GJS32_07500 | 1580104 | 1580376 | -1 | PqqD family peptide modification chaperone |
| QGM14280.1 |  | GJS32_07505 | 1580473 | 1581336 | -1 | helix-turn-helix domain-containing protein |
| QGM14281.1 |  | GJS32_07510 | 1581588 | 1581878 | -1 | hypothetical protein |
| QGM14282.1 |  | GJS32_07515 | 1581894 | 1582193 | -1 | hypothetical protein |
|  |  | GJS32_07520 | 1582264 | 1583637 | -1 | helicase |
|  |  | GJS32_07525 | 1583745 | 1585523 | -1 | reverse transcriptase |
|  |  | GJS32_07530 | 1586033 | 1591498 | -1 | methyltransferase domain-containing protein |
| QGM14283.1 |  | GJS32_07535 | 1591549 | 1592100 | -1 | hypothetical protein |
|  |  | GJS32_07540 | 1592118 | 1594697 | -1 | glucan-binding protein |
| QGM14284.1 |  | GJS32_07545 | 1594857 | 1595480 | -1 | hypothetical protein |
| QGM14285.1 |  | GJS32_07550 | 1595610 | 1596869 | 1 | ISL3-like element ISSeq1 family transposase |
| QGM14286.1 |  | GJS32_07555 | 1597252 | 1597572 | -1 | CHAP domain-containing protein |
| QGM14287.1 |  | GJS32_07560 | 1597527 | 1598369 | -1 | nucleotidyl transferase AbiEii/AbiGii toxin family protein |
| QGM14288.1 |  | GJS32_07565 | 1598366 | 1598959 | -1 | transcriptional regulator |
|  |  | GJS32_07570 | 1599081 | 1601883 | -1 | CHAP domain-containing protein |
| QGM14289.1 |  | GJS32_07575 | 1601880 | 1602848 | -1 | type IV secretion system protein VirB4 |
| QGM14290.1 | ltrA | GJS32_07580 | 1602853 | 1604526 | -1 | group II intron reverse transcriptase/maturase |
| QGM14291.1 |  | GJS32_07585 | 1605055 | 1606542 | -1 | type VI secretion protein |
| QGM14292.1 |  | GJS32_07590 | 1606499 | 1606852 | -1 | PrgI family protein |
| QGM14293.1 |  | GJS32_07595 | 1606914 | 1607768 | -1 | conjugal transfer protein TrbL |
| QGM14294.1 |  | GJS32_07600 | 1607785 | 1608027 | -1 | hypothetical protein |
| QGM14295.1 |  | GJS32_07605 | 1608045 | 1608668 | -1 | TraM recognition domain-containing protein |
| QGM14296.1 | ltrA | GJS32_07610 | 1608691 | 1610601 | -1 | group II intron reverse transcriptase/maturase |
|  |  | GJS32_07615 | 1611410 | 1612684 | -1 | type IV secretory system conjugative DNA transfer family protein |
|  |  | GJS32_07620 | 1612684 | 1613158 | -1 | hypothetical protein |
| QGM14297.1 |  | GJS32_07625 | 1613236 | 1613817 | -1 | CPBP family intramembrane metalloprotease |
| QGM14298.1 |  | GJS32_07630 | 1613820 | 1614053 | -1 | transcriptional regulator |
| QGM14299.1 |  | GJS32_07635 | 1614063 | 1614446 | -1 | hypothetical protein |
| QGM14300.1 |  | GJS32_07640 | 1614456 | 1614884 | -1 | hypothetical protein |
|  | dcm | GJS32_07645 | 1614868 | 1616223 | -1 | DNA (cytosine-5-)-methyltransferase |
| QGM14301.1 |  | GJS32_07650 | 1616378 | 1617232 | -1 | replication initiator protein |
| QGM14302.1 |  | GJS32_07655 | 1617213 | 1617401 | -1 | hypothetical protein |
| QGM14855.1 |  | GJS32_07660 | 1617541 | 1617720 | -1 | DNA mismatch repair protein MutT |
| QGM14303.1 |  | GJS32_07665 | 1618216 | 1619547 | -1 | GntP family permease |
| QGM14304.1 |  | GJS32_07670 | 1619639 | 1620412 | -1 | 3-hydroxybutyrate dehydrogenase |
| QGM14305.1 |  | GJS32_07675 | 1620449 | 1621093 | -1 | 3-oxoacid CoA-transferase subunit B |
| QGM14306.1 |  | GJS32_07680 | 1621099 | 1621749 | -1 | 3-oxoacid CoA-transferase subunit A |
| QGM14307.1 |  | GJS32_07685 | 1621764 | 1622954 | -1 | acetyl-CoA C-acyltransferase |
| QGM14308.1 |  | GJS32_07690 | 1623146 | 1624048 | 1 | LysR family transcriptional regulator |
| GI-5 | | | | | | |
| QGM14546.1 |  | GJS32_09050 | 1897996 | 1898184 | -1 | 50S ribosomal protein L28 |
| QGM14869.1 |  | GJS32_09055 | 1898630 | 1899928 | -1 | ISLre2 family transposase |
| QGM14547.1 |  | GJS32_09060 | 1900081 | 1900380 | -1 | hypothetical protein |
| QGM14548.1 |  | GJS32_09065 | 1900392 | 1901642 | -1 | DNA modification methylase |
| QGM14549.1 |  | GJS32_09070 | 1901654 | 1903297 | -1 | relaxase/mobilization nuclease domain-containing protein |
| QGM14550.1 | mobC | GJS32_09075 | 1903260 | 1903643 | -1 | plasmid mobilization relaxosome protein MobC |
| QGM14551.1 |  | GJS32_09080 | 1903664 | 1903873 | -1 | hypothetical protein |
| QGM14552.1 |  | GJS32_09085 | 1903987 | 1904286 | -1 | hypothetical protein |
| QGM14553.1 |  | GJS32_09090 | 1904338 | 1907577 | -1 | hypothetical protein |
| QGM14554.1 |  | GJS32_09095 | 1907616 | 1909661 | -1 | TraM recognition domain-containing protein |
| QGM14555.1 |  | GJS32_09100 | 1909661 | 1910152 | -1 | DUF3801 domain-containing protein |
| QGM14556.1 |  | GJS32_09105 | 1910174 | 1910956 | -1 | hypothetical protein |
| QGM14557.1 |  | GJS32_09110 | 1910956 | 1911228 | -1 | hypothetical protein |
| QGM14558.1 |  | GJS32_09115 | 1911250 | 1911852 | -1 | hypothetical protein |
| QGM14559.1 |  | GJS32_09120 | 1911876 | 1914572 | -1 | CHAP domain-containing protein |
| QGM14560.1 |  | GJS32_09125 | 1914605 | 1915135 | 1 | hypothetical protein |
| QGM14561.1 |  | GJS32_09130 | 1915329 | 1917680 | -1 | DUF87 domain-containing protein |
| QGM14562.1 |  | GJS32_09135 | 1917661 | 1918020 | -1 | PrgI family protein |
| GI-6 | | | | | | |
| QGM14596.1 |  | GJS32_09425 | 1973842 | 1974003 | 1 | ISL3 family transposase |
| QGM14597.1 |  | GJS32_09430 | 1974514 | 1975296 | -1 | TIR domain-containing protein |
| QGM14598.1 |  | GJS32_09435 | 1975311 | 1975922 | -1 | ScaI family restriction endonuclease |
| QGM14599.1 |  | GJS32_09440 | 1975915 | 1976160 | -1 | helix-turn-helix domain-containing protein |
| QGM14600.1 |  | GJS32_09445 | 1976259 | 1977110 | 1 | site-specific DNA-methyltransferase |
| QGM14601.1 |  | GJS32_09450 | 1977177 | 1978349 | -1 | hypothetical protein |
| QGM14602.1 |  | GJS32_09455 | 1978712 | 1979029 | 1 | hypothetical protein |
| QGM14603.1 |  | GJS32_09460 | 1979010 | 1979357 | 1 | hypothetical protein |
| QGM14871.1 |  | GJS32_09465 | 1979608 | 1980699 | 1 | cell division protein FtsK |
| QGM14604.1 |  | GJS32_09470 | 1980980 | 1982350 | 1 | AAA family ATPase |
| QGM14605.1 |  | GJS32_09475 | 1982418 | 1982678 | 1 | helix-turn-helix domain-containing protein |
|  |  | GJS32_09480 | 1982746 | 1983961 | 1 | tyrosine-type recombinase/integrase |
| QGM14606.1 | rpsI | GJS32_09485 | 1984075 | 1984467 | -1 | 30S ribosomal protein S9 |
| QGM14607.1 | rplM | GJS32_09490 | 1984487 | 1984933 | -1 | 50S ribosomal protein L13 |
| QGM14608.1 |  | GJS32_09495 | 1985471 | 1986331 | -1 | DegV family EDD domain-containing protein |
| QGM14609.1 |  | GJS32_09500 | 1986523 | 1987041 | -1 | NYN domain-containing protein |
| QGM14596.1 |  | GJS32_09425 | 1973842 | 1974003 | 1 | ISL3 family transposase |
| QGM14597.1 |  | GJS32_09430 | 1974514 | 1975296 | -1 | TIR domain-containing protein |
| QGM14598.1 |  | GJS32_09435 | 1975311 | 1975922 | -1 | ScaI family restriction endonuclease |
| QGM14599.1 |  | GJS32_09440 | 1975915 | 1976160 | -1 | helix-turn-helix domain-containing protein |
| QGM14600.1 |  | GJS32_09445 | 1976259 | 1977110 | 1 | site-specific DNA-methyltransferase |
| QGM14601.1 |  | GJS32_09450 | 1977177 | 1978349 | -1 | hypothetical protein |
| QGM14602.1 |  | GJS32_09455 | 1978712 | 1979029 | 1 | hypothetical protein |
| QGM14603.1 |  | GJS32_09460 | 1979010 | 1979357 | 1 | hypothetical protein |
| QGM14871.1 |  | GJS32_09465 | 1979608 | 1980699 | 1 | cell division protein FtsK |
| QGM14604.1 |  | GJS32_09470 | 1980980 | 1982350 | 1 | AAA family ATPase |
| QGM14605.1 |  | GJS32_09475 | 1982418 | 1982678 | 1 | helix-turn-helix domain-containing protein |
|  |  | GJS32_09480 | 1982746 | 1983961 | 1 | tyrosine-type recombinase/integrase |

**Appendix Table 6**. Prevalence of predicted genomic islands of OH-71905 among other 47 *S. zooepidemicus* isolates

| Proportion of CDS for *S. zooepidemicus* isolates in each GIs predicted in OH-71905 | | | | | | |
| --- | --- | --- | --- | --- | --- | --- |
|  | GI-1 | GI-2 | GI-3 | GI-4 | Gi-5 | GI-6 |
| Total CDSs | 4 | 106 | 26 | 58 | 18 | 12 |
| ISU37775 | 3/4 | 6/106 | 6/26 | 6/58 | 1/18 | 0/12 |
| ISU6659 | 4/4 | 23/106 | 7/26 | 37/58 | 1/18 | 0/12 |
| ISU54485 | 2/4 | 10/106 | 6/26 | 31/58 | 1/18 | 1/12 |
| ISU9714 | 3/4 | 46/106 | 6/26 | 9/58 | 1/18 | 1/12 |
| ISU36185 | 3/4 | 8/106 | 6/26 | 6/58 | 1/18 | 0/12 |
| ISU38408 | 3/4 | 16/106 | 6/26 | 16/58 | 1/18 | 1/12 |
| ISU75596 | 2/4 | 8/106 | 5/26 | 8/58 | 1/18 | 0/12 |
| ISU88977 | 4/4 | 28/106 | 6/26 | 8/58 | 1/18 | 1/12 |
| ISU16140 | 4/4 | 21/106 | 6/26 | 8/58 | 1/18 | 1/12 |
| ISU54026 | 2/4 | 9/106 | 6/26 | 31/58 | 1/18 | 1/12 |
| AZ-45470 | 4/4 | 10/106 | 7/26 | 8/58 | 1/18 | 0/12 |
| IA-61192 | 4/4 | 27/106 | 25/26 | 52/58 | 1/18 | 12/12 |
| TN-74097 | 4/4 | 105/106 | 26/26 | 57/58 | 18/18 | 12/12 |
| NVSLBI19 | 3/4 | 9/106 | 5/26 | 9/58 | 1/18 | 0/12 |
| NVSLTN-LIVER4 | 4/4 | 106/106 | 25/26 | 57/58 | 18/18 | 12/12 |
| NVSLTN-LUNG1 | 4/4 | 105/106 | 26/26 | 58/58 | 18/18 | 12/12 |
| NVSLTN-LUNG2 | 4/4 | 105/106 | 26/26 | 58/58 | 18/18 | 12/12 |
| NVSLTN-LUNG3 | 4/4 | 105/106 | 26/26 | 56/58 | 18/18 | 12/12 |
| NVSLTN-TB1 | 4/4 | 88/106 | 26/26 | 54/58 | 17/18 | 12/12 |
| NVSLTN-TC1 | 3/4 | 103/106 | 26/26 | 57/58 | 18/18 | 12/12 |
| NVSLVA-S2 | 4/4 | 90/106 | 25/26 | 58/58 | 1/18 | 12/12 |
| NVSLVA-S19 | 4/4 | 88/106 | 24/26 | 57/58 | 1/18 | 12/12 |
| NVSLVA-S22 | 4/4 | 87/106 | 25/26 | 56/58 | 1/18 | 12/12 |
| 2329 | 3/4 | 14/106 | 5/26 | 9/58 | 1/18 | 1/12 |
| ATCC 35246 | 4/4 | 90/106 | 26/26 | 58/58 | 18/18 | 12/12 |
| BHS5 | 3/4 | 10/106 | 5/26 | 32/58 | 1/18 | 1/12 |
| CY | 4/4 | 35/106 | 26/26 | 58/58 | 18/18 | 12/12 |
| H70 | 2/4 | 10/106 | 6/26 | 31/58 | 1/18 | 1/12 |
| MGCS10565 | 3/4 | 6/106 | 6/26 | 6/58 | 1/18 | 0/12 |
| NCTC4676 | 3/4 | 11/106 | 6/26 | 31/58 | 1/18 | 1/12 |
| NCTC6176 | 2/4 | 27/106 | 6/26 | 7/58 | 1/18 | 1/12 |
| NCTC6180 | 4/4 | 29/106 | 6/26 | 15/58 | 1/18 | 1/12 |
| NCTC7022 | 2/4 | 6/106 | 6/26 | 33/58 | 11/18 | 0/12 |
| NCTC7023 | 2/4 | 6/106 | 6/26 | 33/58 | 11/18 | 0/12 |
| NCTC11606 | 3/4 | 6/106 | 6/26 | 6/58 | 1/18 | 0/12 |
| NCTC11824 | 4/4 | 12/106 | 6/26 | 6/58 | 1/18 | 0/12 |
| NCTC12090 | 3/4 | 9/106 | 6/26 | 7/58 | 1/18 | 0/12 |
| Sz4is | 3/4 | 7/106 | 5/26 | 7/58 | 1/18 | 0/12 |
| Sz5 | 3/4 | 23/106 | 7/26 | 10/58 | 1/18 | 1/12 |
| Sz12is | 3/4 | 5/106 | 7/26 | 22/58 | 1/18 | 0/12 |
| Sz16 | 3/4 | 6/106 | 5/26 | 30/58 | 1/18 | 1/12 |
| Sz35 | 3/4 | 10/106 | 21/26 | 50/58 | 1/18 | 12/12 |
| Sz57 | 4/4 | 14/106 | 5/26 | 9/58 | 1/18 | 1/12 |
| Sz105 | 3/4 | 29/106 | 6/26 | 9/58 | 1/18 | 1/12 |
| SzAM35 | 1/4 | 12/106 | 4/26 | 30/58 | 1/18 | 1/12 |
| SzAM60 | 3/4 | 16/106 | 5/26 | 8/58 | 1/18 | 0/12 |
| SzS31A1 | 3/4 | 5/106 | 6/26 | 6/58 | 1/18 | 0/12 |

**Appendix Table 7**. Distribution of putative virulence genes for 48 *S. zooepidemicus* isolates

| Isolates | Presence of Putative Virulence Genes | | | | | | | | | | | | | | |
| --- | --- | --- | --- | --- | --- | --- | --- | --- | --- | --- | --- | --- | --- | --- | --- |
|  | szP | mlpZ | szM | BifA | fszF | sdzD | spaZ | speK | speL | speM | szeF | szeL | szeM | szeN | szeP |
| ISU37775 | **+** | **+** | **-** | **-** | **+** | **+** | **+** | **-** | **-** | **-** | **-** | **-** | **-** | **-** | **-** |
| ISU6659 | **+** | **-** | **-** | **-** | **-** | **+** | **-** | **-** | **-** | **-** | **+** | **-** | **-** | **-** | **-** |
| ISU54485 | **+** | **-** | **-** | **-** | **-** | **+** | **-** | **-** | **-** | **-** | **-** | **-** | **-** | **-** | **-** |
| ISU9714 | **+** | **+** | **-** | **-** | **-** | **+** | **-** | **-** | **-** | **-** | **-** | **-** | **-** | **-** | **-** |
| ISU36185 | **+** | **-** | **-** | **-** | **-** | **+** | **-** | **-** | **-** | **-** | **-** | **-** | **-** | **+** | **+** |
| ISU38408 | **+** | **-** | **-** | **-** | **+** | **+** | **-** | **-** | **-** | **-** | **+** | **-** | **-** | **-** | **-** |
| ISU75596 | **+** | **-** | **-** | **-** | **-** | **+** | **-** | **-** | **-** | **-** | **-** | **-** | **-** | **+** | **-** |
| ISU88977 | **+** | **+** | **-** | **-** | **-** | **+** | **+** | **-** | **-** | **-** | **-** | **-** | **-** | **-** | **-** |
| ISU16140 | **+** | **+** | **-** | **-** | **-** | **+** | **+** | **-** | **-** | **-** | **-** | **-** | **-** | **-** | **-** |
| ISU54026 | **+** | **-** | **+** | **-** | **-** | **+** | **-** | **-** | **-** | **-** | **-** | **-** | **-** | **-** | **-** |
| AZ-45470 | **+** | **-** | **-** | **-** | **-** | **-** | **-** | **-** | **-** | **-** | **-** | **-** | **-** | **-** | **-** |
| IA-61192 | **+** | **-** | **-** | **+** | **+** | **+** | **+** | **-** | **-** | **-** | **-** | **-** | **-** | **-** | **-** |
| OH-71905 | **+** | **-** | **+** | **+** | **+** | **+** | **+** | **-** | **-** | **-** | **-** | **-** | **-** | **-** | **-** |
| TN-74097 | **+** | **-** | **+** | **+** | **+** | **+** | **+** | **-** | **-** | **-** | **-** | **-** | **-** | **-** | **-** |
| NVSLBI19 | **+** | **+** | **-** | **-** | **-** | **-** | **-** | **-** | **-** | **-** | **-** | **-** | **-** | **+** | **-** |
| NVSLTN-LIVER4 | **+** | **-** | **+** | **+** | **+** | **+** | **+** | **-** | **-** | **-** | **-** | **-** | **-** | **-** | **-** |
| NVSLTN-LUNG1 | **+** | **-** | **+** | **+** | **+** | **+** | **+** | **-** | **-** | **-** | **-** | **-** | **-** | **-** | **-** |
| NVSLTN-LUNG2 | **+** | **-** | **+** | **+** | **+** | **+** | **+** | **-** | **-** | **-** | **-** | **-** | **-** | **-** | **-** |
| NVSLTN-LUNG3 | **+** | **-** | **+** | **+** | **+** | **+** | **+** | **-** | **-** | **-** | **-** | **-** | **-** | **-** | **-** |
| NVSLTN-TB1 | **+** | **-** | **+** | **+** | **+** | **+** | **+** | **-** | **-** | **-** | **-** | **-** | **-** | **-** | **-** |
| NVSLTN-TC1 | **+** | **-** | **+** | **+** | **+** | **+** | **+** | **-** | **-** | **-** | **-** | **-** | **-** | **-** | **-** |
| NVSLVA-S2 | **+** | **-** | **-** | **+** | **+** | **+** | **+** | **-** | **-** | **-** | **-** | **-** | **-** | **-** | **-** |
| NVSLVA-S19 | **+** | **-** | **-** | **+** | **+** | **+** | **+** | **-** | **-** | **-** | **-** | **-** | **-** | **-** | **-** |
| NVSLVA-S22 | **+** | **-** | **-** | **+** | **+** | **+** | **+** | **-** | **-** | **-** | **-** | **-** | **-** | **-** | **-** |
| 2329 | **+** | **-** | **-** | **-** | **-** | **+** | **-** | **-** | **-** | **-** | **+** | **-** | **-** | **-** | **-** |
| ATCC 35246 | **+** | **-** | **+** | **+** | **+** | **+** | **+** | **-** | **-** | **-** | **-** | **-** | **-** | **-** | **-** |
| BHS5 | **+** | **-** | **-** | **-** | **-** | **-** | **-** | **-** | **-** | **-** | **+** | **-** | **-** | **+** | **+** |
| CY | **+** | **-** | **+** | **+** | **+** | **+** | **+** | **-** | **-** | **-** | **-** | **-** | **-** | **-** | **-** |
| H70 | **+** | **-** | **+** | **-** | **-** | **+** | **-** | **-** | **-** | **-** | **-** | **-** | **-** | **-** | **-** |
| MGCS10565 | **+** | **+** | **-** | **-** | **+** | **+** | **+** | **-** | **-** | **-** | **-** | **-** | **-** | **-** | **-** |
| NCTC4676 | **+** | **-** | **-** | **-** | **-** | **+** | **+** | **-** | **-** | **-** | **-** | **-** | **-** | **-** | **-** |
| NCTC6176 | **+** | **-** | **-** | **-** | **-** | **+** | **-** | **-** | **-** | **-** | **+** | **-** | **-** | **-** | **-** |
| NCTC6180 | **+** | **-** | **-** | **-** | **-** | **-** | **-** | **-** | **-** | **-** | **+** | **-** | **-** | **-** | **-** |
| NCTC7022 | **+** | **-** | **-** | **-** | **-** | **-** | **-** | **-** | **-** | **-** | **+** | **-** | **-** | **-** | **-** |
| NCTC7023 | **+** | **-** | **-** | **-** | **-** | **-** | **-** | **-** | **-** | **-** | **+** | **-** | **-** | **-** | **-** |
| NCTC11606 | **+** | **+** | **-** | **-** | **+** | **+** | **+** | **-** | **-** | **-** | **-** | **-** | **-** | **-** | **-** |
| NCTC11824 | **+** | **+** | **-** | **-** | **-** | **-** | **-** | **-** | **-** | **-** | **-** | **-** | **-** | **-** | **-** |
| NCTC12090 | **+** | **-** | **-** | **-** | **-** | **+** | **-** | **-** | **-** | **-** | **-** | **-** | **-** | **+** | **+** |
| Sz4is | **+** | **-** | **-** | **-** | **+** | **-** | **-** | **-** | **-** | **-** | **-** | **-** | **-** | **-** | **-** |
| Sz5 | **+** | **-** | **-** | **-** | **-** | **-** | **-** | **+** | **+** | **+** | **-** | **+** | **+** | **-** | **-** |
| Sz12is | **+** | **-** | **-** | **-** | **+** | **-** | **-** | **-** | **-** | **-** | **-** | **-** | **-** | **-** | **-** |
| Sz16 | **+** | **+** | **-** | **-** | **+** | **-** | **-** | **+** | **+** | **+** | **+** | **+** | **+** | **-** | **-** |
| Sz35 | **+** | **+** | **-** | **+** | **+** | **-** | **-** | **-** | **-** | **-** | **+** | **-** | **-** | **-** | **-** |
| Sz57 | **+** | **-** | **-** | **-** | **-** | **+** | **-** | **-** | **-** | **-** | **+** | **-** | **-** | **-** | **-** |
| Sz105 | **+** | **-** | **-** | **-** | **-** | **+** | **-** | **-** | **-** | **-** | **-** | **-** | **-** | **-** | **-** |
| SzAM35 | **+** | **-** | **-** | **-** | **+** | **-** | **-** | **-** | **-** | **-** | **-** | **-** | **-** | **-** | **-** |
| SzAM60 | **+** | **-** | **-** | **-** | **-** | **-** | **-** | **-** | **-** | **-** | **-** | **-** | **-** | **-** | **-** |
| SzS31A1 | **+** | **-** | **-** | **-** | **+** | **-** | **-** | **-** | **-** | **-** | **-** | **-** | **-** | **-** | **-** |
